# Chiral growth of adherent filopodia

**DOI:** 10.1101/2022.06.09.495577

**Authors:** Wenhong Li, Wen-Lu Chung, Michael M. Kozlov, Ohad Medalia, Benjamin Geiger, Alexander D. Bershadsky

## Abstract

Adherent filopodia are elongated finger-like membrane protrusions, extending from the edges of diverse cell types and participating in cell adhesion, spreading, migration and environmental sensing. The formation and elongation of filopodia are driven by the polymerization of parallel actin filaments, comprising the filopodia cytoskeletal core. Here, we report that adherent filopodia, formed during the spreading of cultured cells on galectin-8-coated substrates, tend to change the direction of their extension in a chiral fashion, acquiring a left-bent shape. Cryo-electron tomography examination indicated that turning of the filopodia tip to the left is accompanied by the displacement of the actin core bundle to the right of the filopodia midline. Reduction of the adhesion to galectin-8 by treatment with thiodigalactoside abolished the filopodia chirality. By modulating the expression of a variety of actin-associated filopodia proteins, we identified myosin-X and formin DAAM1 as major filopodia chirality promoting factors. Formin mDia1, actin filament elongation factor VASP, and actin filament crosslinker fascin were also shown to be involved. Thus, the simple actin cytoskeleton of filopodia, together with a small number of associated proteins are sufficient to drive a complex navigation process, manifested by the development of left-right asymmetry in these cellular protrusions.

## Introduction

The development of left-right asymmetry, directed by the actin cytoskeleton, was observed in individual cells and cell collectives *in vitro* (Chen et al., 2012; Tee et al., 2015; Wan et al., 2011), as well as in a wide variety of *in vivo* developmental systems, such as *Caenorhabditis. elegans* oocytes (Middelkoop et al., 2021; Naganathan et al., 2014), *Drosophila* hindgut and genitalia (Hozumi et al., 2006; Lebreton et al., 2018; Spéder et al., 2006; Taniguchi et al., 2011), and in snails, presenting a classic example of biological chirality (Abe and Kuroda, 2019; Davison et al., 2016; Kuroda et al., 2016). Altogether these data suggest that specific actin filament structures, based on actin-associated proteins, such as diverse formins (DAAM1, Cyk1, snail Dia1) and myosins (1D and 1C), can translate and extend the asymmetric helical structure of actin filaments into asymmetric cellular and multicellular morphologies. The mechanisms underlying these multi-scale transformations, and the roles of individual actin associated proteins in these processes, are still poorly characterized.

Filopodia are relatively simple cylindrical membrane protrusions, the formation of which is driven by actin filament self-organization and membrane remodeling. Specifically, filopodia growth is thought to be driven by actin polymerization at the tip, which is regulated by members of the formin family, mainly mDia2 and FMNL2 (Dimchev et al., 2021; Mellor, 2010), in concert with VASP-family proteins (Barzik et al., 2014; Cheng and Mullins, 2020). It was also shown that the Arp2/3 complex can participate in filopodia initiation at the cell’ s leading edge (Yang and Svitkina, 2011) and that the actin crosslinking protein fascin is required for the straight growth of filopodia (Machesky and Li, 2010; Vignjevic et al., 2006), in cooperation with another formin, DAAM1 (Jaiswal et al., 2013). Several myosins, were shown to be localized in distinct domains of filopodia (Houdusse and Titus, 2021), primarily, myosin-X (myo10), which is enriched at the filopodia tips (Berg and Cheney, 2002; Bohil et al., 2006). The overexpression of myosin-X strongly promotes filopodia formation in many cell types (Kerber and Cheney, 2011). The inverse bar domain protein, IRSp53, is also enriched at the filopodia tips and shafts, and facilitates filopodia formation by inducing negative membrane curvature, and by recruiting several actin associated proteins promoting the assembly of actin filament core (Ahmed et al., 2010; Zhao et al., 2011). Finally, filopodia tips were shown to be adhesive sites, enriched with transmembrane adhesion receptors, such as integrins, as well as the proteins connecting these receptors with the actin core, such as talin (Gallop, 2020; Lagarrigue et al., 2015). Myosin-X cooperates with talin in integrin activation at filopodia tip (Miihkinen et al., 2020).

Filopodia take active roles in cell navigation, migration and invasion, through their functions as environmental sensors (Heckman and Plummer, 2013), that can recognize not only the biochemical properties of the extracellular matrix (ECM) but also its physical properties (Alieva et al., 2019; Wong et al., 2014). In relation to chirality development, it was proposed that filopodia of nerve cell growth cone can direct a large scale left-right asymmetric axonal growth (Tamada et al., 2010).

In the course of a previous study (Li et al., 2021b), we have shown that early spreading of Hela-JW cells on substrate coated with the animal lectin, galectin-8, is characterized by an intense formation of filopodia, extending radially from the cells’ edge, while being tightly attached to the substrate, as can be inferred from interference reflection microscopy images. Based on these initial observations, we further revealed that galectin-8 adherent filopodia can deviate from the common pattern of straight, radial extension, and change their growth direction in a defined left-right asymmetric manner. We investigated the dynamic and structural changes underlying the asymmetric behavior of this seemingly simple actin-filament based structure. We show here that the asymmetric turning of filopodia depends on their adhesion to the substrate, that myosin-X and formin DAAM1 are the primary drivers of turning chirality, and that the actin polymerization regulators mDia1, mDia2, and VASP, as well as the actin crosslinking protein fascin can fine-tune the development of filopodia chirality.

## Results

### Filopodia turn counter-clockwise in cells spreading on galectin-8

The vast majority (>90%) of filopodia, extended by Hela-JW cells, during their spreading on galectin-8-coated substrate, are largely straight, while a small fraction is bent (mostly in a counter-clockwise fashion) (Fig 1A). Overexpression of myosin-X or knock-down of Arp2, a key component of the Arp2/3 complex, increased the length of filopodia at twenty minutes following cell plating on galectin-8, without affecting the filopodia number (Fig 1B, 1C). This increase in length was accompanied by a striking increase in the fraction of bent filopodia (Fig 1A, D). Further analysis of these bent filopodia revealed an apparent left-right asymmetry (chirality) in their shape (Fig 1A, F), manifested by a major prevalence of left-bent filopodia.

**Figure 1:**
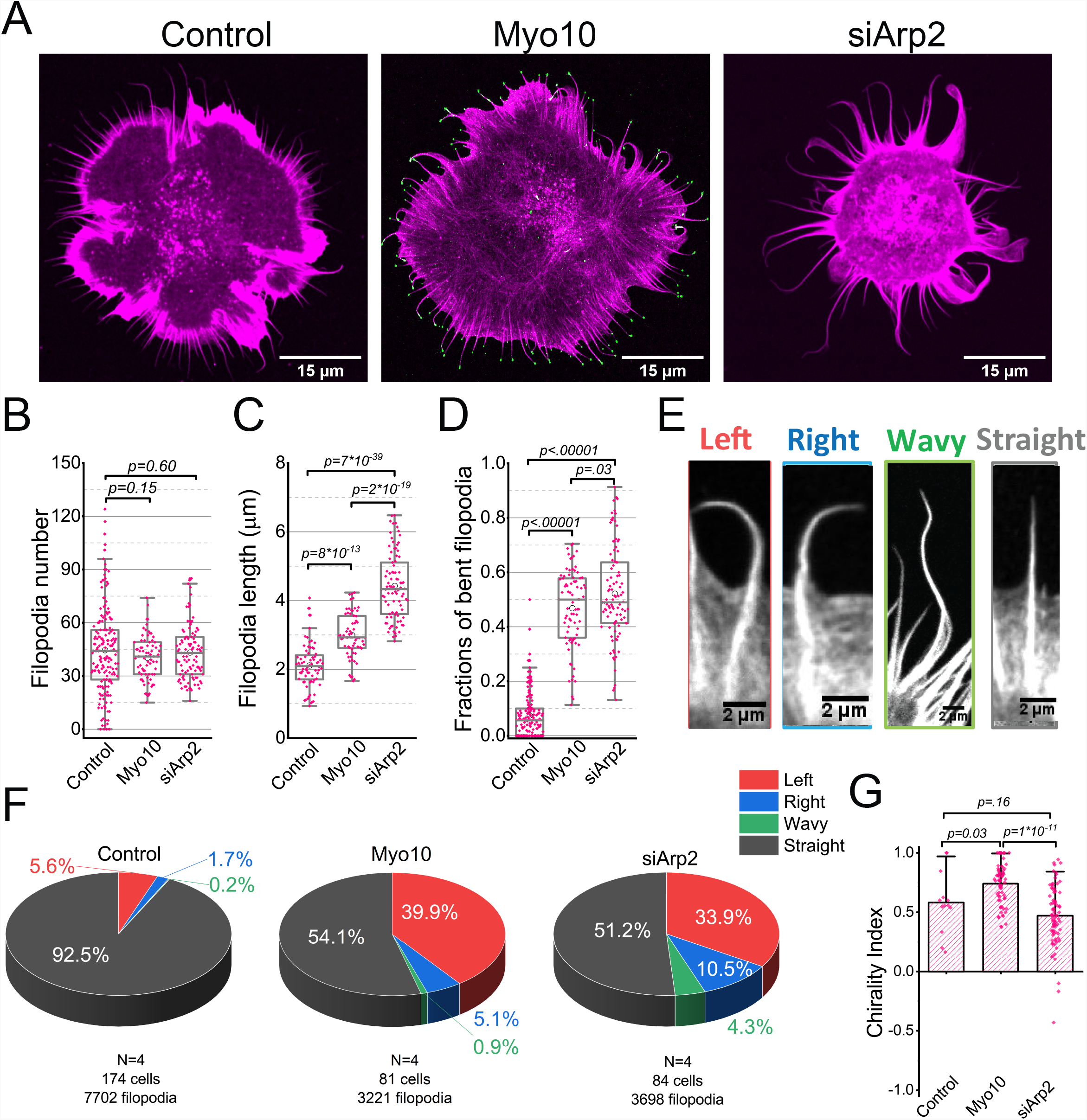
Filopodia turn counter-clockwise in cells spreading on galectin-8. **A**. Confocal images of typical HeLa-JW cells, non-transfected (left), transfected with GFP-Myosin-X (middle), and treated with siRNA against Arp2 (right). Cells were fixed 20 minutes following spreading on galectin-8 coated substrate and stained with phalloidin to visualize actin (magenta). GFP-Myosin-X is shown in green. **B-D**. Filopodia number (B), length (C), and fractions of bent filopodia (D) in HeLa-JW cells, non-transfected (control), transfected with GFP-Myosin-X (Myo10), and treated with siRNA against Arp2 (siArp2), as assessed 20 minutes following plating on galectin-8 coated substrate. Each dot corresponds to an individual cell. Pooled data from more than three experiments in each case are presented as box & whisker plots. *p* values were calculated using non-parametric Mann-Whitney test. **E**. Typical images of left-bent, right-bent, wavy, or straight filopodia. **F**. The pie diagrams representing percentage of the filopodia with different morphology in cells treated as indicated. Sectors of the pies, corresponding to left-bent, right-bent, wavy filopodia and straight filopodia are denoted by red, blue, green and gray, respectively. Data on non-transfected (control), GFP-myosin-X transfected (Myo10), and Arp2 knockdown (siArp2) cells were calculated by assessment of all filopodia from pooled experiments. The number of experiments (N), number of cells assessed, and total number of filopodia are indicated under each pie. **G**. Chirality index is defined as the number of left-bent minus right bent filopodia divided by the total number of non-straight filopodia. Each dot corresponds to an individual cell. Pooled data from no less than three experiments were presented. The bars show the mean values and error bars the standard deviation. *p* values were calculated using non-parametric Mann-Whitney test.

To assess this asymmetry, we calculated, for each cell, a chirality index, defined as (L-R)/(L+R+W) where L, R and W are the numbers of the left-bent, right-bent, and “wavy” filopodia respectively. The “wavy filopodia” are those filopodia that change direction more than once, along their length (Fig 1E), and are observed relatively rarely (Fig 1F) and thus, they were not included in the chirality index numerator. To further clarify the nature of the chirality index: if all curvy filopodia are bent left, the chirality index is equal to 1, and if all curvy filopodia are bent right, the chirality index would be -1. An index equals to 0 corresponds to an identical number of left and right bent filopodia (lack of chirality).

In control, non-transfected cells, spreading on galectin-8 the fraction of bent filopodia was low (7.4% ± 8.5%, N=174 cells), but these few filopodia clearly demonstrated asymmetric bending with a chirality index of 0.58± 0.07. Arp2 knockdown resulted in many more bent filopodia, the majority of which turned left. This effect was observed upon Arp2 knockdown using two different single siRNAs (Fig S2). The chirality index of filopodia following Arp2 knockdown did not, however, exceed the values observed in control cells (Fig 1G). Myosin-X overexpression also increased the fraction of bent filopodia, and in addition significantly augmented their chirality, resulting in a chirality index of up to 0.76± 0.02. Overexpression of truncated mutants of myosin-X, lacking the FERM domain (Myo10-ΔFERM) and even lacking the entire complex of C-terminal PH, MyTH4 and FERM domains (Myo10-HMM) also increased the fraction of bent filopodia and, remarkably, their chirality index (Fig S3). A similar increase in filopodia chirality was observed using an additional cell line, mouse melanoma B16F1 (Fig S1), pointing to the generality of the chiral turning phenomenon.

Time lapse imaging of myosin-X expressing cells and Arp2 knockdown cells, spreading on galectin-8, clearly show that the “bent” morphology of filopodia occurs upon a change in the direction of filopodia growth, rather than by physical bending of existing, fully extended filopodia (Fig 2A, Movie 1, 2), and the alteration of the direction of filopodia growth is often preceded by a decrease in filopodia growth velocity (Fig 2B). In cells expressing GFP-myosin-X, the change of filopodium direction is often accompanied by an apparent duplication of myosin-X positive patch at the tip of filopodium. The GFP-myosin-X patch associated with the “old” tip stops moving while the new, tilted segment of the filopodium, acquire a new GFP-myosin-X patch at its tip (Fig 2C). In less than 20 seconds, however, the old myosin-X patch disappears and the new filopodium front segment becomes curved (Fig 2E, Movie 3).

**Figure 2:**
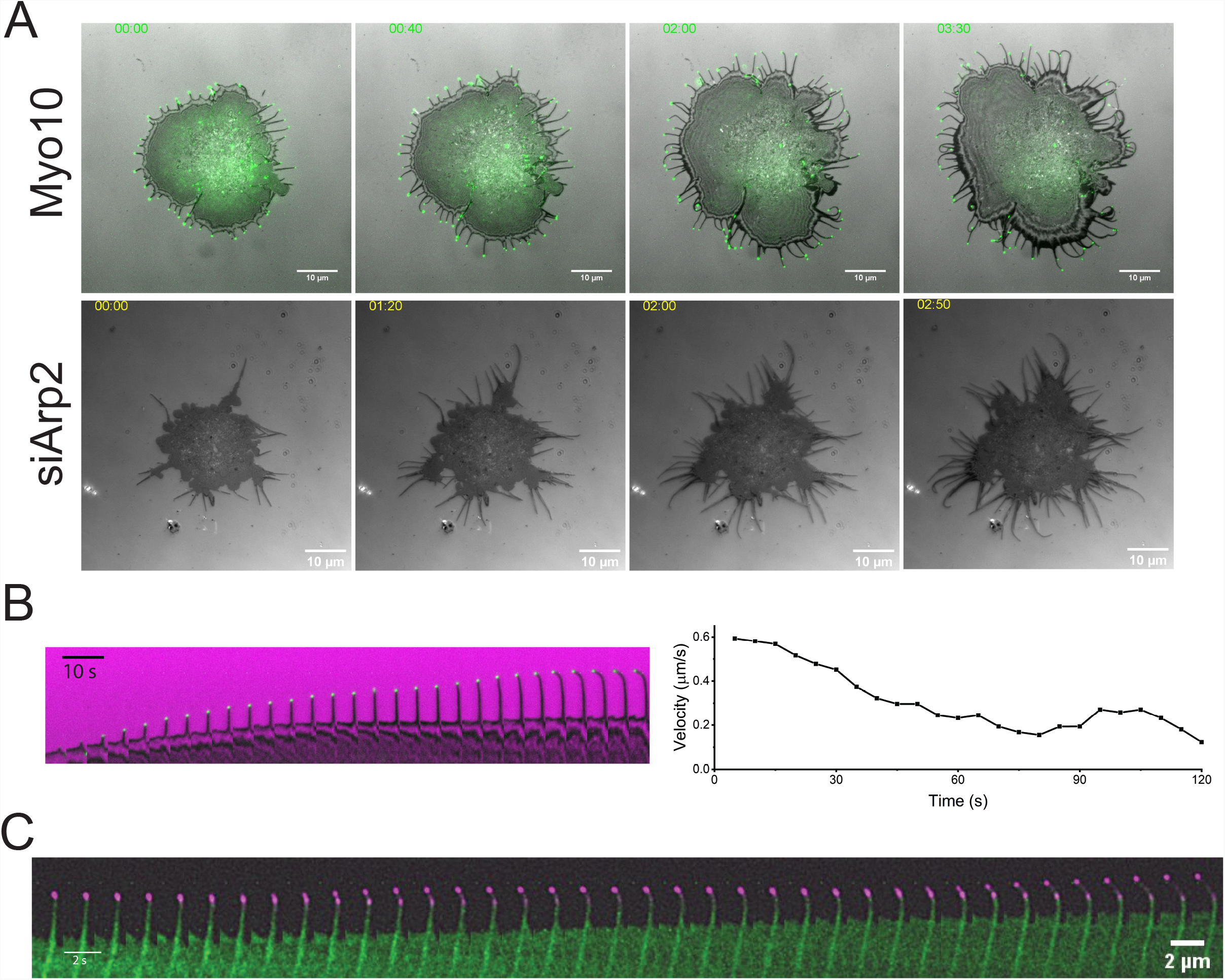
Dynamics and structure of turning filopodia. **A**. Time course of the formation of bent filopodia by changing the direction of extension of straight filopodia in cells transfected with GFP-Myosin-X (Myo10) or Arp2 knockdown cells (siArp2) during cell spreading on substrates coated with galectin-8. Cells were imaged using interference reflection microscopy (IRM), in combination with epi-fluorescence microscopy for visualization of GFP-Myosin-X. Scale bars: 15 µm. Time after cell seeding (minutes) is indicated in each image. **B**. Left: Kymograph showing the time course of extension and turning of a typical filopodium of cell transfected with GFP-Myosin-X. Notice the bent filopodium was formed from a straight filopodium which changed the direction of extension. Right: The velocity of the filopodia tip relative to the substrate during the growth period calculated for the filopodia shown in the left. Notice that the velocity of filopodia tip is reduced before filopodia change the direction of extension. **C**. Kymograph showing the myosin-X (magenta) dynamics during filopodia turning process. Note the filopodia turning is accompanied by splitting of the myosin-X patch into two. One is associated with the former end of the filopodia as it was before turning and eventually disappeared, the other is associated with the new filopodia tip.

The results of Cryo-EM examination of filopodia induced by overexpression of GFP-myosin-X are presented in Fig 3. The straight and bent filopodia can be distinguished at lower magnification (insights in tomograms A-F). Higher magnification of filopodia tips revealed that in many cases, the cylindrical filopodia were ended by bulbous extensions. Among bent filopodia, almost half (47%) had such bulb-like ending while straight filopodia displayed such structure in much rarer cases (5.6%).

**Figure 3:**
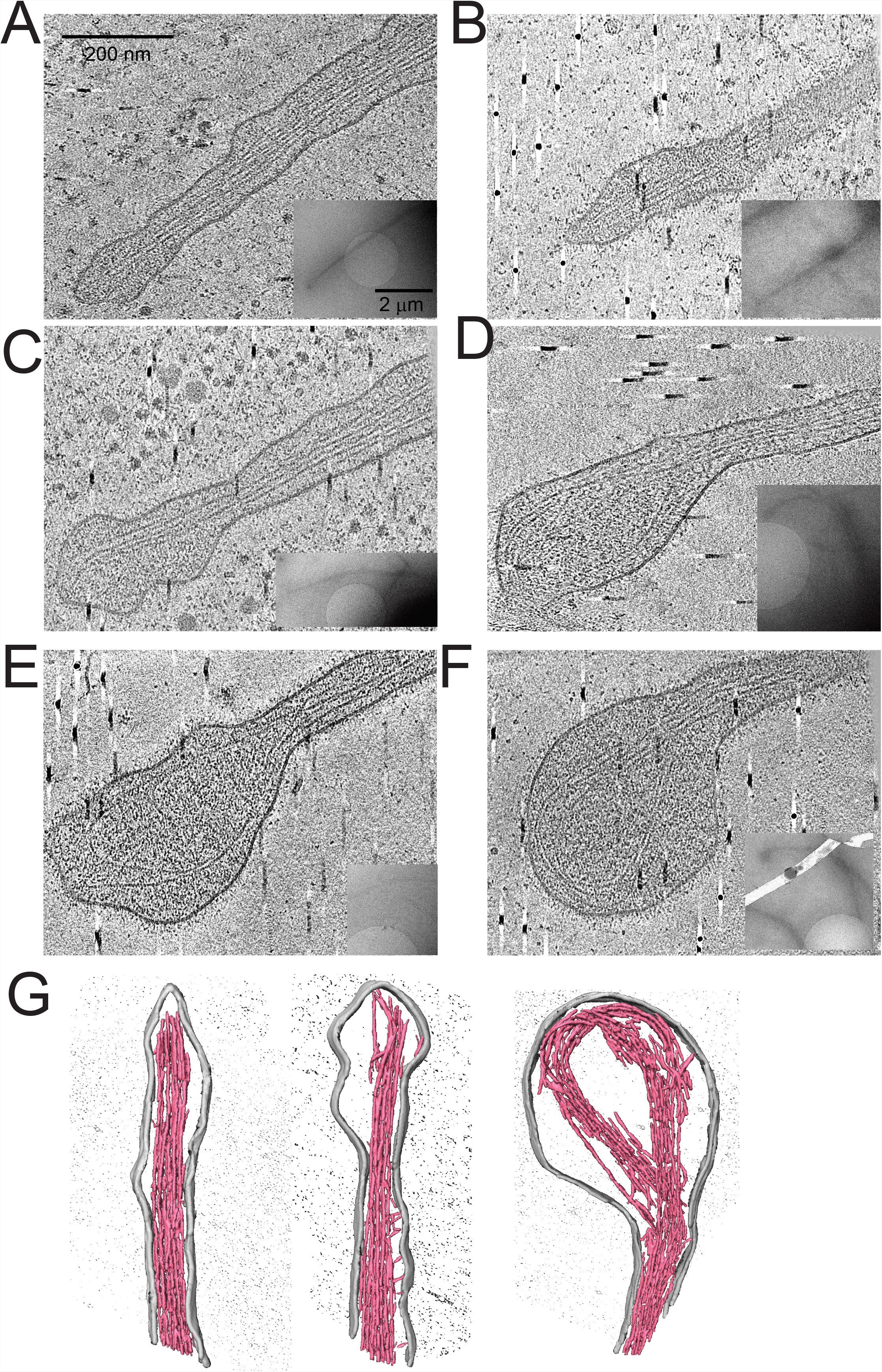
Cryo-electron tomography images of filopodia tips. **A-F**. A slice of Cryo-ET images in x-y plane of straight filopodia (**A** and **B**) and left bent filopodia (**C**-**F**) in cells overexpressing GFP-Myosin-X spreading on galectin-8 coated substrates for 20 minutes. Slice thickness: 8.9 nm, see the movies 4-9 respectively. **G**. 3D rendering isosurface of the images in **B, C**, and **F** (red: actin, gray: membrane). Note that in straight filopodia (A, B and G-left) the filament cores are located symmetrically along the mid-line of filopodia and approached the filopodia tips. In the bent filopodia (C-F, G-middle and right), the bulbous membrane extensions are located asymmetrically relatively to the bulk of the actin cores, which are shifted to the right side of the bulbs. The left halves of the bulbs either did not contain actin filaments (C, G-middle), or contained splayed filaments (D, E), or the loop formed by the continuation of the core (F, G-right).

The filopodia actin core appeared as a tight bundle of essentially continuous filaments (Fig 3). Short filament fragments found sometimes in the filopodia (Fig 3A-F, also shown in 3D rendering isosurface, Fig 3G) are expected due to the resolution limitation in the condensed environment. In straight filopodia, the actin core was located symmetrically in the centre of the membrane tube (Fig 3A, B, G), while in bent filopodia, the position of the actin core relative to the membrane was always asymmetric at the filopodia tip (Fig 3C-F, G). The segment of the actin core bundle in the proximity of filopodium tip was always located at the side opposite to the bulb, so that the distance from the core to right edge of filopodium was always smaller than to the left edge (Fig 3C-F, G). Thus, in the majority of filopodia tips, the membrane bulbs were shifted to the left and either did not contain actin filaments at all (Fig 3C) or contained few filaments splayed from the actin bundles (Fig 3D, E). However, in some cases, the actin bundles aligned along the right edge of the tip, bent to the left and sometimes form a loop along the edge of bulb (Fig 3F, G).

### Adhesion to galectin-8 coated substrate is essential for chiral turning

Interference reflection microscopy (IRM) shows that filopodia of cells, growing on galectin-8 coated substrates, maintain a close proximity to the substrate (IRM-dark areas in Fig 2 and (Li et al., 2021b). To determine whether the tight adhesion to the underlying matrix is needed for the chiral turning, we added to the spreading cells, thiodigalactoside (TDG), a homolog of galectins ligand β-galactoside, which is known to block the binding of galectin-8 to its membrane receptors (Levy et al., 2001; Li et al., 2021). As shown in Fig 4, TDG reduces filopodia formation, though numerous filopodia were still formed and the fraction of bent filopodia was slightly reduced in the presence of 10 mM TDG (Fig 4A, B). Remarkably, however, TDG reduced chirality of filopodia in both myosin-X expressing and Arp2 depleted cells, so that the fractions of left and right turning filopodia were approximately equal (Fig 4A, B).

**Figure 4:**
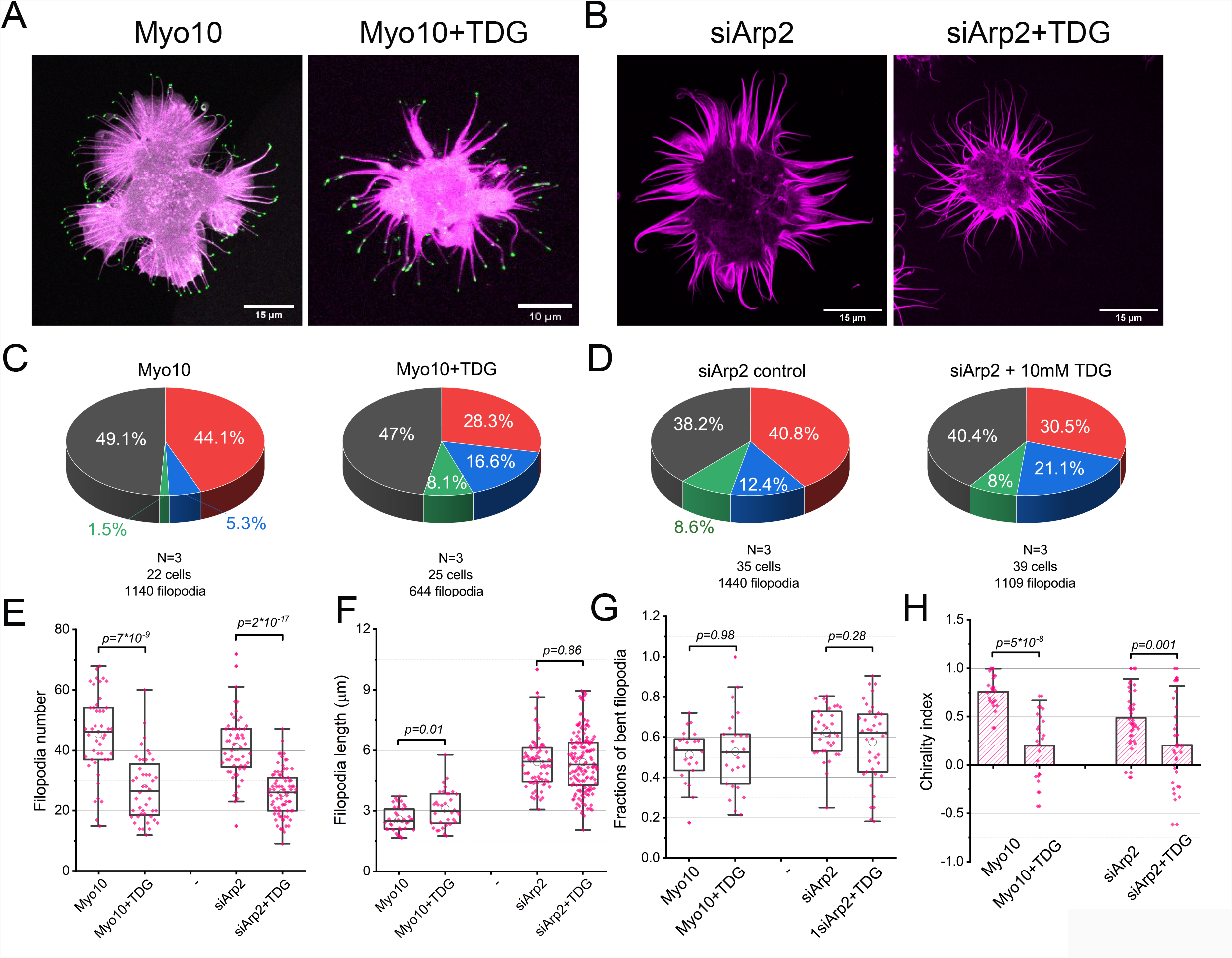
Reduction of cell adhesion to galectin-8 by thiodigalactoside abolished filopodia chirality. **A.B**. Confocal images of HeLa-JW cells expressing GFP-Myosin-X (Myo10) (A) and depleted of Arp2 (siArp2) (B) spreading on galectin-8 coated substrate for 20 minutes in the absence or in the presence of 10mM galectin-8 ligand thiodigalactoside (TDG). Cells were stained with phalloidin to visualize actin (magenta). GFP-Myosin-X is shown in green. **C.D**. The pie diagrams representing percentage of the filopodia with different morphology in cells treated as indicated. Color coding is the same as in Figure 1. The number of experiments (N), cells and filopodia assessed are indicated under each pie. Note that incubation with TDG significantly decreased the fraction of left-bent filopodia (C) and increased the fraction of right-bent filopodia (C, D). **E-G**. Filopodia number (E), length (F), and fractions of bent filopodia (G) in cells treated as indicated. Each dot corresponds to an individual cell. Pooled data from two experiments for each condition are presented as box & whisker plots. *p* values were calculated using non-parametric Mann-Whitney test. **H**. Filopodia chirality index in cells treated as indicated. Each dot corresponds to an individual cell. Pooled data from two experiments were presented. The bars show the mean values and error bars the standard deviation. *p* values were calculated using non-parametric Mann-Whitney test. Note that treatment with TDG reduced filopodia chirality.

### Filopodia chiral turning is myosin-X-dependent

Myosin-X was shown to be important for filopodia formation (Bohil et al., 2006), and consequently, myosin-X depleted cells form significantly fewer filopodia than control cells, even on galectin-8 coated substrates (Fig S3G, S4B). Since the fraction of bent filopodia was rather low in both control and myosin-X knockdown cells (Fig S3F, G, S4A, B) and given that Arp2 knockdown strongly enhances the fraction of bent filopodia (Fig 1), we first checked whether myosin-X function is required for asymmetric filopodia turning after Arp2 knockdown. As shown in Figure 4, double depletion of Arp2 and myosin-X indeed resulted in the formation of larger number of bent filopodia than the knockdown of myosin-X alone. Among these bent filopodia, the fractions of left and right bent filopodia were approximately equal (Fig 5A, B). This result was observed upon knockdown of myosin-X by Dharmacon™ smartpool siRNAs as well as by the individual siRNAs constituting the smartpool (Fig S4D-F). Thus, depletion of myosin-X strongly decreases filopodia chirality in Arp2 knock-down cells while, myosin-X overexpression increases filopodia chirality in Arp2 knockdown cells (Fig 5).

**Figure 5:**
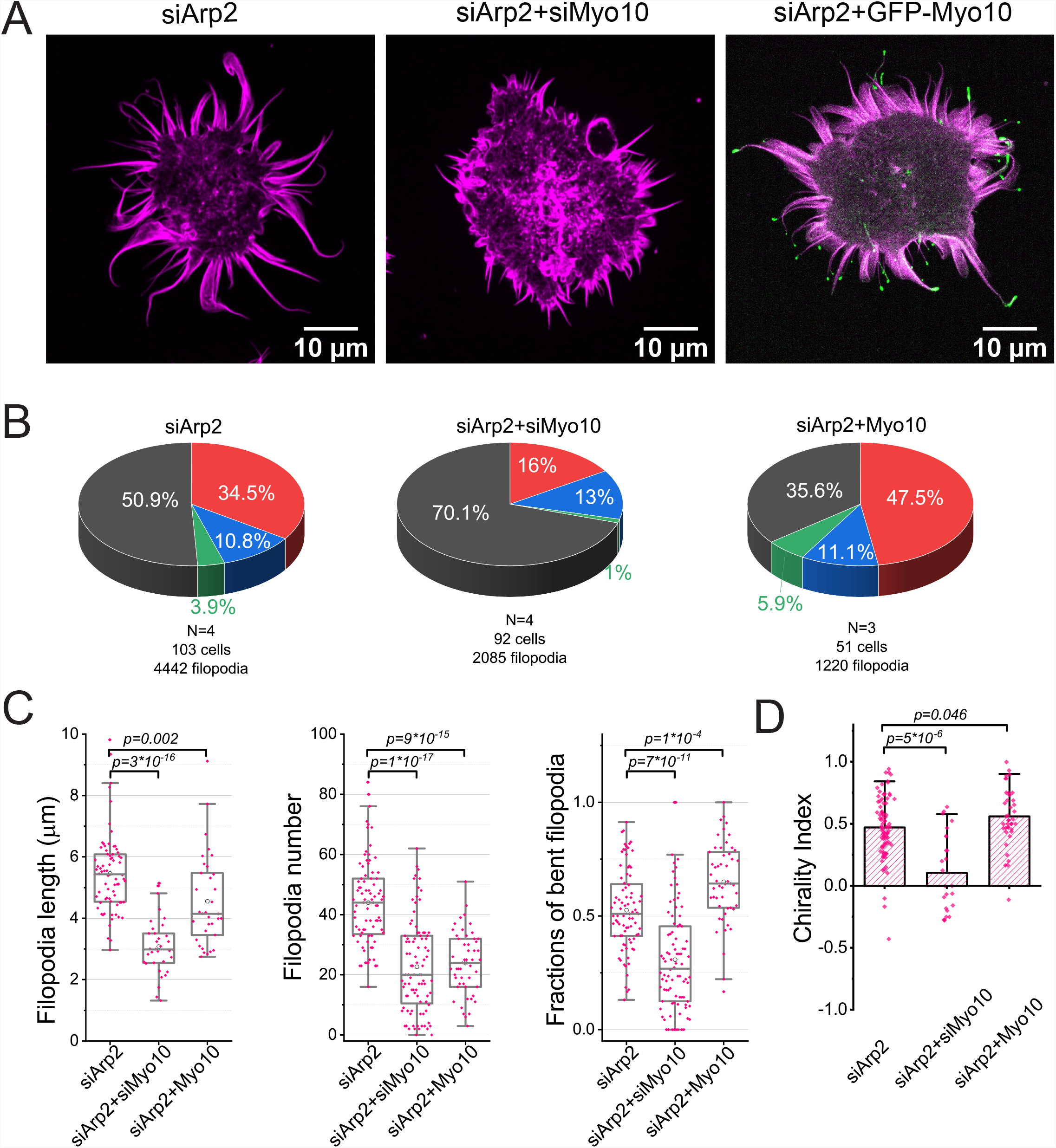
Myosin-X is required for filopodia chirality in Arp2 knockdown cells. **A**. Confocal images of Arp2 depleted cells (siArp2), Arp2 and myosin-X depleted cells (siArp2+siMyo10), and Arp2 depleted cells expressing GFP-Myosin-X (siArp2+GFP-Myo10) fixed 20 minutes following spreading on galectin-8 coated substrate and stained with phalloidin to visualize actin (magenta). GFP-Myosin-X is shown in green. **B**. The pie diagrams representing percentage of the filopodia with different morphology in cells treated as indicated. Color coding is the same as in Figure 1. The numbers of experiments (N), cells and filopodia assessed are indicated under each pie. Note that myosin-X knockdown significantly reduced the fraction of left-bent filopodia while its overexpression increased that fraction. **C**. Filopodia number, length, and fractions of bent filopodia in cells treated as indicated. Each dot corresponds to an individual cell. Pooled data from two experiments for each condition are presented as box & whisker plots. *p* values were calculated using non-parametric Mann-Whitney test. **D**. Filopodia chirality index in cells treated as indicated. Each dot corresponds to an individual cell. Pooled data from two experiments were presented. The bars show the mean values and error bars the standard deviation. p values were calculated using non-parametric Mann-Whitney test. Note that knockdown of myosin-X reduced filopodia chirality index.

We further show that filopodia turning chirality following myosin-X knockdown, can be restored in human Hela-JW cells, by the expression of bovine (rather than the human) protein (Fig S3). This significantly increased the number of filopodia, their length and their filopodia turning chirality. Interestingly, truncated myosin-X mutants also rescued the formation and chirality of filopodia in myosin-X knockdown cells. Filopodia formed by cells lacking endogenous myosin-X and expressing tailless myosin-X-HMM instead, remained chiral even though to a less extent than filopodia expressing full length myosin-X (Fig S3). Chirality of filopodia in cells expressing myosin-X lacking FERM domain (GFP-Myo10-ΔFERM) was like that in control cells (Fig S3).

We further checked whether myosin-X induces chirality also in cells with unperturbed Arp2 levels. Given the very low number of bent filopodia in control and myosin-X knockdown cells, we have tested many cells, enabling us to reliably assess and compare the chirality index of their filopodia. Our results showed that knockdown of myosin-X significantly decreases the chirality of filopodia in cells spreading on galectin-8 (Fig S4). These results, together with the data shown in Fig 1b, indicate that myosin-X is essential for the development of filopodia chirality, irrespective of Arp2 levels.

### Myosin-1D and 1C depletion does not affect filopodia turning chirality

Several other members of the myosin super-family were previously shown to be involved in different forms of cell chirality. Myosin 1D and 1C were shown to be critical for the development of chirality in *Drosophila* embryos (Hozumi et al., 2006; Lebreton et al., 2018; Spéder et al., 2006). Myosin 1C binds to phosphatidylinositol 4, 5-bisphosphate (PIP_2_) and was shown to be clustered at the tips as well as the base of a special type of membrane extension, stereocilia (García et al., 1998; Gillespie and Müller, 2009).

As shown in Fig. S5, both myosin 1D and 1C affect filopodia formation in control and myosin-X overexpressing cells; myosin 1D apparently down-regulates filopodia formation (based on the observation that its knockdown enhances filopodia number and length, while its overexpression suppresses filopodia), and myosin 1C promoted filopodia formation (its knockdown reduced filopodia number in control cells and filopodia length in myosin-X overexpressing cells). However, despite these effects, neither myosin 1D nor myosin 1C depletion affected the fraction of bent filopodia or filopodia bending chirality (Fig S5).

### Effects of depletion of regulators of actin polymerization at the filopodia tip on chiral turning

Filopodia growth is driven by actin polymerization at the tip of filopodia, which is regulated, locally, by formin and VASP family proteins. Among the major formins in Hela cells (Krainer et al., 2013), mDia1, mDia2 and FMNL2 were reported to be enriched at the filopodia tips (Block et al., 2012; Goh and Ahmed, 2012; Mellor, 2010; Yang et al., 2007). VASP was also found to be localized at the tip of filopodia (Applewhite et al., 2007). Here, we directly investigated the functions of mDia1, mDia2 and FMNL2, as well as VASP, in the regulation of filopodia chiral turning on galectin-8. We examined how depletion of these molecules affect filopodia growth in control cells and cells where enhanced formation of bent filopodia was induced by myosin-X overexpression or Arp2 knockdown.

mDia2 depletion strongly reduced filopodia number and length in control cells, but overexpression of myosin-X or knockdown of Arp2 significantly rescued filopodia formation and elongation in cells without mDia2 (Fig S5). Remarkably, depletion of mDia2 significantly increased the chirality index of filopodia in cells overexpressing myosin-X or lacking Arp2 (Fig 6). Of note, changes in chirality index in these experiments did not correlate with the fraction of bent filopodia, since upon mDia2 knockdown, this fraction decreased in Arp2 knockdown cells but increased in myosin-X expressing cells (Fig 5B). Similarly, mDia2 and FMNL2 knockdown also moderately increased filopodia chirality in Arp2 knockdown cells but did not affect filopodia chirality in cells overexpressing myosin-X (Fig 6). Thus, the two formins that play a major role in filopodia growth, mDia2 and FMNL2, appear to be dispensable for filopodia chirality and may even suppress chiral turning.

**Figure 6:**
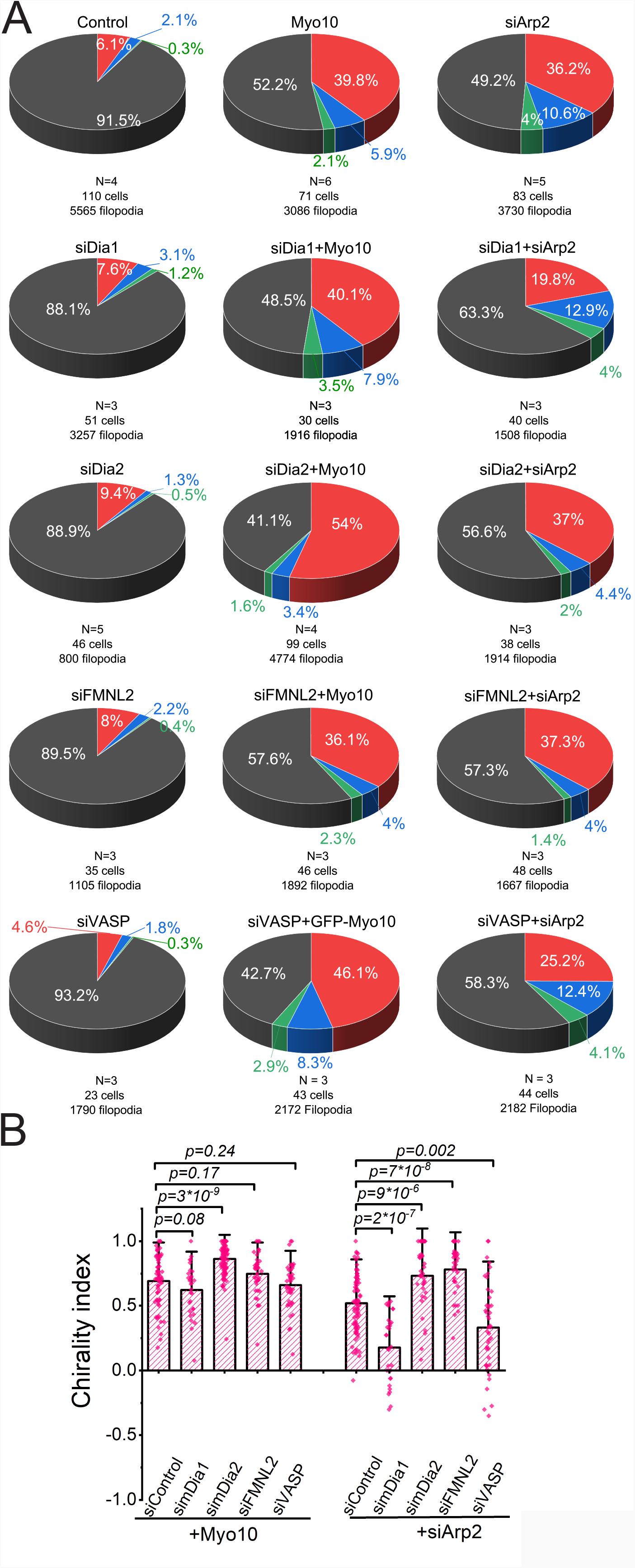
The effects of knockdowns of mDia1, mDia2, FMNL2, and VASP on filopodia chirality. **A**. The pie diagrams representing percentage of the filopodia with different morphology in cells treated as indicated. Color coding is the same as in Figure 1. The numbers of experiments (N), cells and filopodia assessed are indicated under each pie. In the left column, the data for non-transfected (control) cells and cells with single knockdown of mDia1, mDia2 FMNL2, or VASP are presented. The middle column shows the data for GFP-myosin-X expressing cells (Myo10), without or with the knockdown of mDia1, mDia2 FMNL2, or VASP, respectively. In the right column, the data for Arp2 depleted cells without or with additional knockdown of mDia1, mDia2 FMNL2, or VASP are shown. **B**. Filopodia chirality index in cells treated as indicated. Each dot corresponds to an individual cell. Pooled data from two experiments were presented. The bars show the mean values and error bars the standard deviation. *p* values were calculated using non-parametric Mann-Whitney test. Note that depletion of mDia1 and VASP reduced chirality in Arp2 knockdown cells, while depletion of mDia2 and FMNL2 slightly increased filopodia chirality index.

Knockdown of mDia1 slightly increased filopodia number in control and myosin-X overexpressing cell but not in Arp2 depleted cells, and had, essentially, no effects on filopodia length (Fig 6). It also did not affect the fraction of curved filopodia and the chirality index in control and myosin-X overexpressing cells (Fig 5). However, in Arp2 knockdown cells, depletion of mDia1 resulted in a pronounced decrease in both the fraction of bent filopodia, and the filopodia chirality index (Fig 6, S5). Thus, while mDia1 is dispensable for filopodia chirality in control and myosin-X overexpressing cells, filopodia chiral turning in Arp2 depleted cells requires mDia1. VASP overexpression was shown to reduce filopodia formation in Hela cells spreading on galectin-8 coated substrate (Li et al., 2021b). Like mDia1, VASP depletion significantly reduced the fractions of curved filopodia and filopodia chirality index in cells lacking Arp2. Thus, filopodia chiral turning in Arp2 depleted cells requires myosin-X, mDia1, and VASP.

### Requirement of filopodia shaft proteins DAAM1, and fascin for chiral turning of filopodia

We next set to investigate several proteins that are mainly located at the filopodia shaft. Actin filament crosslinking protein fascin along with formin DAAM1 are reported to function to bundle actin filaments (Jaiswal et al., 2013). In addition, DAAM1 is one of the highly expressed formins in Hela cells (Krainer et al., 2013). Thus, we examined the roles of fascin and DAAM1 in regulating filopodia chirality.

The depletion of DAAM1 significantly increased filopodia length and the fraction of bent filopodia in control as well as in myosin-X overexpressing and Arp2 depleted cells (Fig S7). The fraction of wavy filopodia also increased upon DAAM1 depletion (Fig S7). Strikingly, however, DAAM1 depletion resulted in the loss of filopodia chirality (Fig 7). The depletion of fascin dramatically inhibited filopodia number and length. It also increased the fractions of bent filopodia in control and myosin-X overexpressing cells but not in Arp2 knockdown cells. Fascin depletion significantly decreased the chirality in myosin-X overexpressing cells, even though the chirality index remained positive. In Arp2 knockdown cells as well as in control cells, the fascin depletion did not reduce the ratio between left and right turning filopodia and therefore did not reduce the chirality index. However, the fraction of wavy filopodia was increased markedly by fascin knockdown in these cases too, resulting in less chiral phenotype. Thus, while both DAAM1 and fascin are required for filopodia chirality, the effect of DAAM1 depletion was more pronounced than that of fascin depletion.

**Figure 7:**
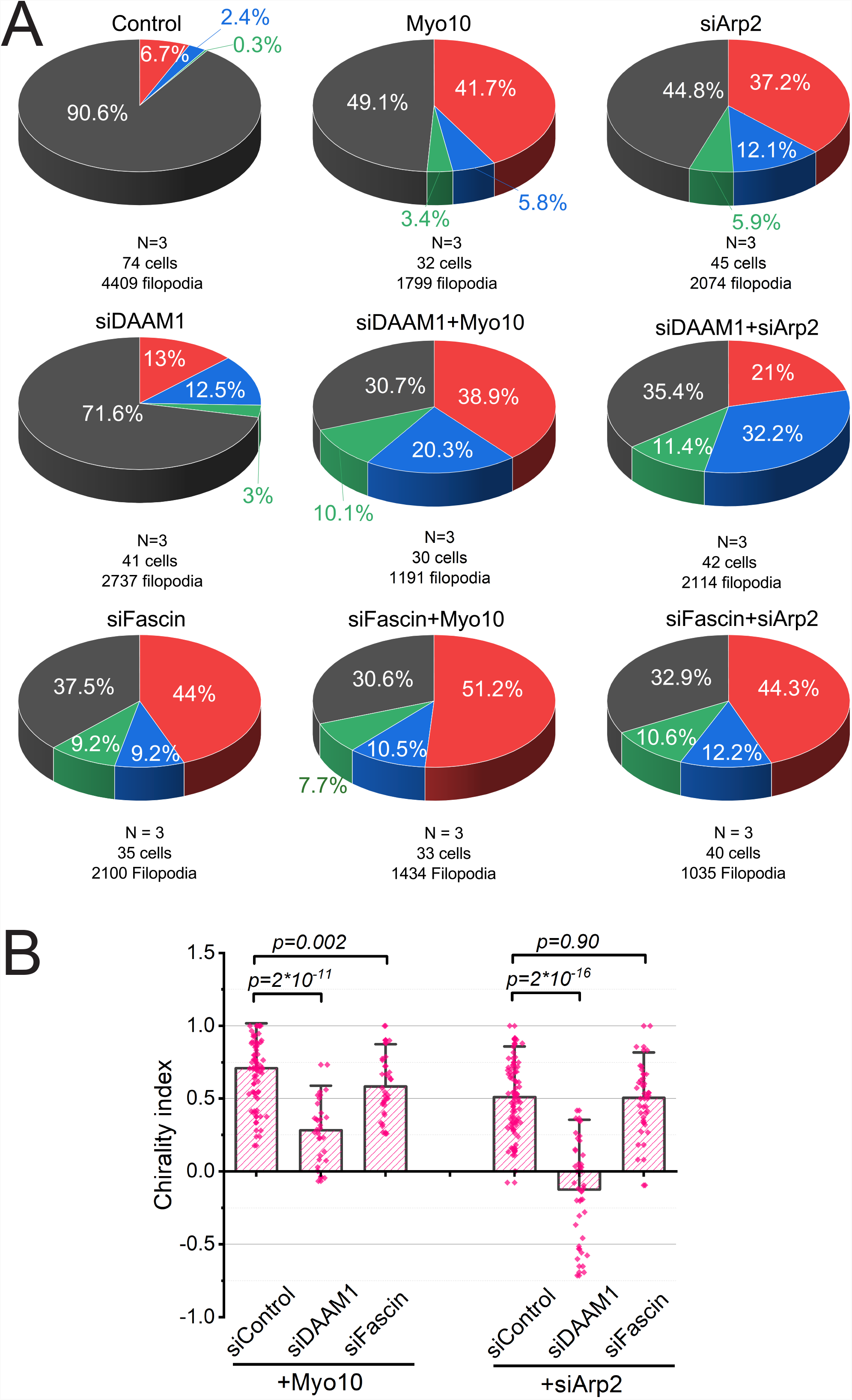
The effects of knockdown of fascin and DAAM1 in filopodia formation and filopodia chirality. **A**. The pie diagrams representing percentage of the filopodia with different morphology in cells treated as indicated. Color coding is the same as in Figure 1. The numbers of experiments (N), cells and filopodia assessed are indicated under each pie. In the left column, the data for non-transfected (control) cells and cells with single knockdown of DAAM1 or fascin are presented. The middle column shows the data for GFP-myosin-X expressing cells (Myo10), without or with the knockdown of DAAM1 or fascin, respectively. In the right column, the data for Arp2 depleted cells without or with additional knockdown of DAAM1 or fascin are shown. Note that knockdown of DAAM1 increased the fraction of right-bent filopodia while knockdown of fascin reduced the fraction of straight filopodia and increased the fraction of right-bent and wavy filopodia. **B**. Filopodia chirality index in cells treated as indicated. Each dot corresponds to an individual cell. Pooled data from two experiments were presented. The bars show the mean values and error bars the standard deviations. *p* values were calculated using non-parametric Mann-Whitney test. Note that knockdown of DAAM1 reduced chirality index stronger than knockdown of fascin.

## Discussion

In this study, we observed a new phenomenon: a chiral turning of matrix-associated filopodia during their elongation. We showed here that growing filopodia, attached to a planar substrate, uniformly coated with galectin-8, tend to turn, more often, to the left than to the right, during their growth. An asymmetric movement of filopodia was previously observed in some nerve cells, where a straight filopodium moves as a generatrix of a cone with its apex coinciding with the filopodium base and the tip following a circular path in one direction (Tamada and Igarashi, 2017; Tamada et al., 2010). Unlike these non-adherent filopodia rotations, the chirality of filopodia in our system emerges due to an asymmetric change in the direction of filopodium extension, resulting in the formation of filopodia that are curved in the plane of the substrate. To address the mechanisms underlying this asymmetric turning, three major aspects of filopodia growth were considered: the elongation dynamics, the involvement of the adhesion to extracellular matrix, and the cytoskeletal machinery that, most likely, regulates the turning direction.

Even though the galectin-8 coated substrate stimulates the formation of adherent filopodia, the vast majority of them remain straight and only few display a chiral turning. We found two main conditions that strongly increase the fraction of turning filopodia: (i) overexpression of myosin-X or (ii) knockdown of Arp2 expression, thereby suppressing Arp2/3 mediated actin polymerization. Under both conditions, filopodia length increases and the changes in filopodia growth direction occur in 40%-60% of filopodia. The majority of these filopodia turned left (counter-clockwise), as seen by an observer looking at the cell “top-down”, from the medium towards the substrate. These initial observations enabled a systematic study of the molecular mechanism underlying the process of filopodia chiral extension.

Previous studies revealed that adherent filopodia are growing in an intermittent fashion, with periods of extension that are alternating with the period of growth cessation and formation of tighter adhesions between filopodia tips and the substrates (Watanabe et al., 2010). We have shown here that changes in the direction of filopodia extension occur after such intermittent cessation of growth when the new segment of filopodium start to extend from the adherent tip. This suggests the possibility that filopodia adhesion to the substrate is essential for the chiral turning of the filopodia tip. To address this possibility, we have tested the effect of the galectin-8 inhibitory ligand, TDG, on the filopodia turning chirality. As shown in Fig 3, the inhibitor indeed abolished the chiral growth of the filopodia, confirming that adhesion to galectin-8 is needed for the induction of turning chirality.

Searching for the cytoskeletal protein(s) that induce, or support filopodia chirality, we used a systematic siRNA mediated down regulation approach for identifying cytoskeletal components that affect the turning chirality without blocking filopodia growth. Fortunately, the formation of filopodia and their elongation appear to be highly robust processes, so that diverse combinations of the actin associated proteins could be sufficient to promote filopodia growth and the absence of some of them can be compensated (Dobramysl et al., 2021). This is consistent with our results, showing that knockdown of myosin-X blocks the formation of filopodia on galectin-8 coated substrate, but combination of this knockdown with knockdown of Arp2 rescued filopodia formation, and permitted us to obtain enough bent filopodia and assess their chirality. This experiment clearly demonstrated that myosin-X is required for chiral filopodia turning even under conditions when it is dispensable for filopodia growth. Moreover, overexpression of myosin-X not only increases the fraction of bent filopodia but significantly increases their chirality. Myosin-X function as a filopodia chirality inducer is consistent with known characteristics of its molecular motor activity. Myosin-X is moving processively along actin filament bundles that are crosslinked by fascin (Caporizzo et al., 2018; Ropars et al., 2016) in a chiral helical fashion (Sun et al., 2010). Even though the function of myosin-X in filopodia is thought to be related to delivery of different cargos to the filopodia tips (Tokuo and Ikebe, 2004), our experiments show that its involvement in filopodia chirality depends mainly on its interaction with the actin core. The truncated mutants of myosin-X, lacking the C-terminal domains involved in cargo binding, still support filopodia chirality, and myosin-X lacking the FERM domain induces chirality even better than the wild type protein.

Besides myosins, the formin family proteins constitute another class of molecules that can, in principle, create asymmetric torque in the actin filament systems. Specifically, during formin driven actin polymerization, a mobile formin dimer, at the tip (barbed end) of the growing actin filaments, is expected to rotate upon incorporation of each new actin subunit, following the helical structure of the actin filament (Mizuno et al., 2011). When such rotation is restricted (e.g., by formin immobilization), the polymerizing actin filament should rotate in a chiral fashion, which may underlie the development of large scale left-right asymmetry, affecting the self-organization of the actin cytoskeleton in the cell (Tee et al., 2021; Tee et al., 2015). Since formins are localized to filopodia and play a critical role in their growth, we examined the involvement of some of the filopodial formins in the generation of filopodia chirality.

mDia2 (a product of DIAPH3 gene) is one of the highly expressed formins localized to filopodia tips. mDia2 was shown to be required for the formation of filopodia, induced by a the small Rho GTPase Rif (Pellegrin and Mellor, 2005). This is consistent with our initial demonstration that knockdown of mDia2 suppresses the formation of filopodia in Hela-JW cells on galectin-8 coated substrate (Li et al., 2021b). We show here that the formation of filopodia in mDia2-depleted cells can be rescued by overexpression of myosin-X or knockdown of Arp2. These filopodia were chiral with chirality index even higher than that observed in filopodia induced by overexpression of myosin-X or knockdown of Arp2 in mDia2-containing cells. Another major formin localizes to the filopodia tips, FMNL2, was also dispensable for filopodia chiral turning and its knockdown even slightly increases chirality in filopodia of Arp2 knockdown cells. Thus, both major promoters of filopodia growth, mDia2 and FMNL2, are not required for filopodia chirality or even function as its negative regulator. It is interesting to note that actin filament rotation that could be generated by immobilized formins at the filopodia tip should occur in a direction that is opposite to the one generated by myosin-X, and therefore could interfere with it.

Unlike mDia2 and FMNL2, the depletion of another diaphanous family formin, DAAM1, significantly suppressed the filopodia chirality. This effect was especially prominent in cells where filopodia formation was stimulated by myosin-X overexpression or Arp2 knockdown, but can also be observed in non-stimulated cells, spreading on galectin-8 coated substrate. DAAM1 is localized mainly to filopodia shafts due to interaction with the actin cross-linking protein fascin (Jaiswal et al., 2013). In our experiments, the knockdown of fascin also reduced the filopodia chirality. Thus, DAAM1 could promote chirality due to its participation in stabilization of actin core bundles in cooperation with fascin. This suggestion does not explain, however, why the knockdown of DAAM1 suppressed filopodia chirality stronger than the knockdown of fascin.

Two other regulators of actin polymerization were shown here to participate in filopodia chirality regulation. mDia1 is shown to function at the filopodia shafts where it can affect actin polymerization and possibly act as F-actin crosslinker (Goh and Ahmed, 2012), and VASP, which is enriched at filopodia tips, is thought to be a major actin “elongator”, together with the formins mDia2 and FMNL2/3 (Damiano-Guercio et al., 2020). Both mDia1 and VASP, in our experiments, were required for the filopodia chirality in cells depleted of Arp2 but not in cells overexpressing myosin-X. The reason for this specificity is presently unclear.

Thus, our survey of filopodial actin associated proteins revealed that even in this relatively simple system, the development of chirality depends on several players including in particular myosin-X, formin DAAM1, as well as several actin regulators such as mDia1, fascin and VASP. Furthermore, the set of proteins involved in filopodia chiral turning differs from that involved in other types of actin-based chirality. While DAAM1 protein important for chiral morphogenesis in Drosophila developments (Chougule et al., 2020) is indispensable for filopodia chirality, the homologs of other Drosophila chirality proteins, myosin-1C and 1D (Hozumi et al., 2006; Lebreton et al., 2018; Spéder et al., 2006), did not seem to affect the filopodia chirality in our cells. Similarly, the knockdown of mDia1 formin affected filopodia chirality in much less degree than the chiral swirling in human fibroblasts and chiral alignment of these cells (Tee et al., 2021). At the same time, we have found the important function of myosin-X in filopodia chirality, which was not detected in previously studied examples of actin based chiral morphogenesis.

Despite all the differences in particular molecular players, all actin-based chiral systems have apparent common features. They all are based on interactions of helical actin filaments with two classes of proteins, formins and myosins, which can result in the generation of chiral movement. Such chiral movement (for example, axial filament twirling, or twisting of filament bundle) can be a basis for higher order chiral morphogenesis if it is acting in a well-defined system of coordinates. Such coordinate system can be defined by cell adhesion to the extracellular matrix or/and to other cells. In our system, the adhesion of cells and their filopodia to the planar substrate covered with galectin-8 is a necessary condition for the filopodia asymmetric growth.

Taken together, the possible molecular mechanisms underlying filopodia turning chirality, share a common expectation, namely a local structural asymmetry in the organization of the tip of turning filopodia. A direct support to such possibility was provided by observations of turning filopodia using cryo-electron tomography (Fig 3).

The first observation is that in filopodia that start turning left, the actin core bundle is shifted to the right. Thus, two questions should be addressed: (i) why the core actin bundle is shifting and (ii) how this shifting of the core lead to turning of the filopodium tip to the opposite side. Addressing the first question, namely, the shifting of the core, can be explained by its interactions with molecules of myosin-X, which move along a helical path wrapped around the core (Sun et al., 2010). The movement of myosin-X molecules toward the tip of filopodia is accompanied by their counterclockwise rotation around the axis of the core, which may be a reason for the asymmetric accumulation of the myosin-X molecules between the core and the membrane at the left side of the core tip. If this myosin-X rotation is counteracted by some forces such as the friction with the surrounding cytoplasm and/or the interaction with the membrane, the angular momentum will be, partially, transmitted to the core and drive the clockwise axial rotation of the latter. Such core rotation can produce the core “rolling” along the substrate-immobilized membrane to the right until the core distal end gets pushed against the right side of the filopodial membrane. Thus, shifting of the core towards the right edge of filopodia could be a consequence of an accumulation of myosin-X molecules on the left side of the filopodium tip and the myosin-X driven rolling of the core towards the right.

As per the second question, namely how can such shift trigger a leftward turning of the filopodia, several general mechanisms might be considered. Recent published data suggests that filopodia growth is initiated by membrane remodeling with inverted BAR-domain proteins (Jones et al., 2020), while consequent polymerization of actin filaments stabilizes the protrusions and promotes their elongation. Assuming the concentration of the I-BAR domain proteins to be homogeneous all over the filopodial membrane, the mean membrane curvature (½ (c_1_+c_2_) = ½ (1/r_1_+1/r_2_), where c_1_ and c_2_ are the principal curvatures, and r_1_ and r_2_ the principal radii of curvature, respectively) should be constant over the entire filopodia surface. Since in the cylindrical filopodium shaft one of the principal curvatures is zero, the radius of the filopodium’ s spherical tip will be two-fold larger than that of the filopodium’ s cylindrical shaft, i.e. the tip will be swollen compared to the shaft. Indeed, we observed that in many cases the filopodia ended with a bulb with a radius higher than the radius of the shaft. Because of this swelling, the core cannot stick to the membrane of the tip in a symmetrical fashion like in the shaft. In the absence of factors shifting the core such as myosin-X, the core will stick to the right or to the left of the bulb with an equal probability. However, because myosin-X induces systematic shifting of the core to the right, the probability of the core attachment of the right side of the bulb membrane will be higher than probability of the attachment to the left side. It means that the bulb at the tip of filopodia will be shifted to the left relatively to the axis of the shaft. It is reasonable to suggest that this asymmetric bulb can serve as a bud for a newly forming segment of filopodia. Elongation of this segment can be driven by either new actin filament nucleation at the bud, or by partial straightening of the looped actin core.

It should be noted here that presently we are not aware of physiological processes in which left-right asymmetric filopodia extension is involved. Given the role of filopodia in environmental sensing and cellular navigation (Heckman and Plummer, 2013), it might be considered that asymmetric formation of filopodia at some structures, such as growth cone of nerve cells, could partake in the process of chiral growth of neurites, which was discovered many years ago (Heacock and Agranoff, 1977). Filopodia also participate in a variety of processes related to the formation of cell-cell junctions (Li et al., 2021a; Vasioukhin et al., 2000) and cell fusion (Hammers et al., 2021; Segal et al., 2016). For some of these processes, left-right asymmetry could be important, and intrinsic filopodia chirality may underly such asymmetry. Irrespective to its particular role, the mechanism of asymmetric filopodia extension deserves to be further studied. Filopodia provide a challenging example of architecturally simple structures built of actin filaments and membrane, yet displaying new emerging features, such as chiral turning that might affect cellular morphogenesis.

## Acknowledgements

We thank Dr R. E. Cheney (University of North Carolina at Chapel Hill, North Carolina), Dr N. O. Alieva (A-star, Singapore), Dr Yee Han Tee and Dr P. Kanchawong (Mechanobiology Institute, Singapore) for the cells and the constructs. We thank Dr Yaming Jiu (Chinese Academy of Science, China), and Dr Yee Han Tee (Mechanobiology Institute, Singapore) for the discussions on the manuscript.

B.G. is grateful to the Israel Science Foundation (personal grant (2749/17) and precision medicine grant (3617/19)), to the Minerva Center at the Weizmann Institute “Aging, from Physical Materials to Human Tissues” for their support and to the Jean-Jacques Brunschwig Fund for the Molecular Genetics of Cancer for supporting this research.

A.D.B. acknowledge the support from the Singapore Ministry of Education Academic Research Fund Tier 2 (MOE Grant No: MOE2018-T2-2-138), the National Research Foundation, Prime Minister’ s Office, Singapore, and the Ministry of Education under the Research Centers of Excellence program through the Mechanobiology Institute, Singapore (ref no. R-714-006-006-271), and Singapore Ministry of Education Academic Research Fund Tier 3 MOE grants no. MOE2016-T3-1-002, and no. MOET32021-0003.

A.D.B., B.G. and W.L. acknowledge the support from a Maimonides Israeli–France grant (Israeli Ministry of Science Technology and Space), and EU Marie Skłodowska-Curie Network InCeM (project ID 642866).

W.L.C. and O.M. acknowledge ERC-Syg (810057-HighResCells) grant.

---

**Figure S1: Mouse embryo fibroblast and melanoma B16 cells exhibit filopodia chirality upon overexpression of GFP-Myosin-X**

**A and B**. Confocal images of typical mouse embryo fibroblast (A) and melanoma B16 cells (B), non-transfected (left image in each panel) and transfected with GFP-Myosin-X (right image in each panel). Cells were fixed 20 minutes following plating on galectin-8 coated substrate and stained with phalloidin to visualize actin (magenta). GFP-Myosin-X is shown in yellow.

**C and D**. The pie diagrams representing percentage of the filopodia with different morphology in mouse embryo fibroblast cells (C) and melanoma B16 cells (D), non-transfected (left pie in each panel) and transfected with GFP-Myosin-X (right pie in each panel). Color coding is the same as in Figure 1. The number of experiments (N), cells and filopodia assessed are indicated under each pie. Note in both cell types, myosin-X overexpression increased the fraction of bent filopodia and these filopodia demonstrate chirality.

**Figure S2: Validation of the data on effects of Arp2 knockdown on the fractions of bent filopodia and chiral filopodia**

**A**. Confocal images of Hela-JW cells treated with control siRNA (left) and with two individual siRNAs against Arp2 (middle and right panels). Cells were fixed 20 minutes following plating on galectin-8 coated substrate and stained with phalloidin to visualize actin (magenta).

**B**. Pie diagrams representing percentage of the filopodia with different morphology in cells treated as indicated. Color coding is the same as in Figure 1. The numbers of experiments (N), cells and filopodia assessed are indicated under each pie.

**C**. Fractions of bent filopodia in cells treated as indicated. Each dot corresponds to an individual cell. Pooled data from two experiments for each condition are presented as box & whisker plots. p values were calculated using non-parametric Mann-Whitney test.

**D**. Filopodia chirality index in cells treated as indicated. Each dot corresponds to an individual cell. Pooled data from two experiments were presented. The bars show the mean values of chirality indices and error bars -the standard deviation. *p* values were calculated using non-parametric Mann-Whitney test.

**E**. Western blot showing Arp2 protein levels in control siRNA treated cells (control), cells treated with smart pool siRNA against Arp2 (siArp2 smart pool), and cells treated with two individual siRNA against Arp2 (siArp2 siRNA1, and siArp2 siRNA2). The sequences are shown in Materials and Methods.

Note that individual siRNA produces the same effects on filopodia bending and chirality as smartpool siRNA against Arp2. See also Fig 1A, D, F, G.

**Figure S3: Truncated forms of Myosin-X still enhance filopodia chirality**.

**A**. A cartoon depicting the sequences of the full length and truncated forms of myosin-X.

**B-H:** characterization of filopodia after transfection of different construct of myosin-X into control HelaJW cells (B-E) and Hela-JW cells depleted of endogenous myosin-X (F-H).

**B**. Confocal images of HeLa-JW cells, transfected with GFP-Myosin-X (left), GFP-Myosin-X-HMM (middle) and GFP-Myosin-X-ΔFERM (right), fixed 20 minutes following spreading on galectin-8 coated substrate. Actin (magenta) was visualized by phalloidin-Alexa647 staining. GFP-Myosin-X and GFP-Myosin-X mutants are shown in green.

**C and F**. Pie diagrams representing percentage of the filopodia with different morphology 20 minutes following spreading on galectin-8 in Hela-JW cells (C) and in Hela-JW cells transfected with siRNA against myosin-X (F) expressing full length and truncated forms of myosin-X. Color coding is the same as in Figure 1. The numbers of experiments (N), cells and filopodia assessed are indicated under each pie.

**D and G**. The graphs showing (from left to right in each panel), filopodia number, length, and fractions of bent filopodia in Hela-JW cells (D) and the same cells transfected with siRNA against myosin-X (G) treated as indicated. Each dot corresponds to an individual cell. Pooled data from four experiments for each condition are presented as box & whisker plots. *p* values were calculated using non-parametric Mann-Whitney test.

**E and H**. Filopodia chirality indices in Hela-JW cells (E) and the same cells transfected with siRNA against myosin-X (H) treated as indicated. Each dot corresponds to an individual cell. Pooled data from two experiments were presented. The bars show the mean values of chirality indices and error bars - the standard deviation. *p* values were calculated using non-parametric Mann-Whitney test.

Note that, expression of myosin-X lacking FERM domain or containing only the motor domains promoted filopodia chirality in cells both with and without endogenous myosin-X.

**Figure S4: Myosin-1D and myosin-1C are not required for filopodia chirality**.

**A**. Pie diagrams representing percentage of the filopodia with different morphology in Hela-JW cells, non-treated, overexpressing myosin-X, or lacking Arp2, which were in addition transfected with siRNAs against myosin-1D (siMyo1D) and myosin-1C (siMyo1C). Color coding is the same as in Figure 1. The numbers of experiments (N), cells and filopodia assessed are indicated under each pie.

**B-D**. Filopodia number, length, and fractions of bent filopodia in cells treated as indicated. Each dot corresponds to an individual cell. Pooled data from four experiments for each condition are presented as box & whisker plots. *p* values were calculated using non-parametric Mann-Whitney test.

**E**. Filopodia chirality index in cells treated as indicated. Each dot corresponds to an individual cell. The bars show the mean values and error bars the standard deviation (SD). *p* values were calculated using non-parametric Mann-Whitney test.

**F**. Western blot showing myosin-1C and myosin-1D protein levels in control siRNA treated cells (control), and cells treated with smart pool siRNA. Tubulin was used as a loading control.

**Fig S5: Additional data on the effects of knockdowns of mDia1, mDia2, FMNL2, and VASP on filopodia (See figure 5)**

**A-C:** Filopodia number, length, and fractions of bent filopodia in cells treated as indicated. Each dot corresponds to an individual cell. Pooled data from four experiments for each condition are presented as box & whisker plots. *p* values were calculated using non-parametric Mann-Whitney test.

**D:** Western blot showing mDia1, FMNL2 and VASP protein levels in control siRNA treated cells (control), and cells treated with corresponding smart pool siRNAs.

**Fig S6: Additional data on the effects of knockdowns of fascin and DAAM1 on filopodia (See figure 6)**

**A-C:** Filopodia number, length, and fractions of bent filopodia in cells treated as indicated. Each dot corresponds to an individual cell. Pooled data from two experiments for each condition are presented as box & whisker plots. *p* values were calculated using non-parametric Mann-Whitney test.

**D:** Western blot showing fascin and DAAM1 protein levels in control siRNA treated cells (control), and cells treated with corresponding smart pool siRNA.

**Movie 1: Filopodia extension in cells overexpressing myosin-X spreading on galectin-8 coated substrates**

Representative time-lapse series showing a cell transfected with GFP-Myosin-X spreading on the galectin-8 coated substrate, imaged by interference reflection microscopy (IRM) together with fluorescent microscopy (green). The time interval is 5 seconds. Scale bar: 10 μm. Timestamp: mm:ss, The display rate: 3 frames per second.

**Movie 2: Filopodia extension in cells with Arp2 depletion spreading on galectin-8 coated substrates**

Representative time-lapse series showing a cell with Arp2 knockdown spreading on the galectin-8 coated substrate, imaged by interference reflection microscopy (IRM) together with fluorescent microscopy (green). The time interval is 5 seconds. Scale bar: 10 μm. Timestamp: mm:ss, The display rate: 15 frames per second.

**Movie 3: Filopodia extension in cells overexpressing myosin-X spreading on galectin-8 coated substrates**

Representative time-lapse series showing in detail filopodia extension in a cell transfected with GFP-Myosin-X and TD-tomato-F-tractin spreading on the galectin-8 coated substrate, imaged by structured illumination microscopy. The time interval is 5 seconds. Scale bar: 10 μm. Timestamp: mm:ss, The display rate: 7 frames per second.

**Movie 4-5: Cryo-electron tomography of tips of typical straight filopodia**

Slices of Cryo-ET images in x-y plane of straight filopodia in cells overexpressing GFP-Myosin-X spreading on galectin-8 coated substrates for 20 minutes. Slice thickness: 8.9 nm. Tilt angle: from -60° to 60°.

**Movie 6-9: Cryo-electron tomography of tips of typical left-bent filopodia**

Slices of Cryo-ET images in x-y plane of left-bent filopodia in cells overexpressing GFP-Myosin-X spreading on galectin-8 coated substrates for 20 minutes. Slice thickness: 8.9 nm. Tilt angle: from -60° to 60°.

## Materials and Methods

### Cell culture, DNA constructs, and reagents

Hela-JW is a subline of a HeLa cervical carcinoma cell line derived in the laboratory of J. Willams (Carnegie-Mellon University, USA) based on their better attachment to plastic dishes (Bai et al., 1993; Paran et al., 2006). In addition to Hela-JW cells, in some experiments we used primary mouse embryo fibroblast (MEF) at early passage level and mouse melanoma B16 cells (Fidler, 1975). The cells were cultured in Dulbecco’ s modified Eagle’ s medium (DMEM) supplemented with 10% fetal bovine serum (FBS), 1 mM sodium pyruvate and 100 U/mL penicillin-streptomycin in 5% CO2 incubator at 37 °C. The cell culture reagents were purchased from Biological Industries, Ltd. (Beit HaEmek, Israel), and used according to the manufacturer’ s instructions. No cell lines used in this study were found in the database of commonly misidentified cell lines that is maintained by ICLAC and NCBI Bio Sample. We did not attempt to authenticate them.

Cells were transiently transfected with the following DNA plasmids. tdTomato-F-tractin (Schell et al., 2001) was received from Dr. Y. H. Tee (Mechanobiology Institute in Singapore). Three myosin-X constructs, full length myosin-X (1-2052), GFP-Myo10-ΔFERM (1-1916) and GFP-Myo10-HMM (1-843) were produced in the lab of R. E. Cheney (University of North Carolina at Chapel Hill, North Carolina) (Berg and Cheney, 2002; Bohil et al., 2006). GFP-Myo1D was purchased from ADDGENE (#134906). All the transfections were done using JetPEI (Polyplus-transfection® SA, France) following the manufacturer’ s protocols.

### Transfection of siRNA

Cells were seeded into a 35 mm dish on day 0 and transfected with 20 μM siRNA against of Arp2, mDia2, FMNL2, mDia1, DAAM1, VASP, fascin, and myosin-X (Dharmacon, ON-TARGET plus SMART pool siRNA) using Lipofectamine RNAiMAX (ThermoFisher, catalog number: LMRNA015) on day 1 and day 2. Control cells were transfected with scrambled control siRNA, ON-TARGET plus Non-targeting pool siRNA (Dharmacon, catalogue no. D-001810-10). Cells were imaged on day 3 after transfection, 20 minutes following trypsinization and re-plating on galectin-coated substrate. The siRNA sequences in the smart pools were summarized in Table 1.

**Table 1:**
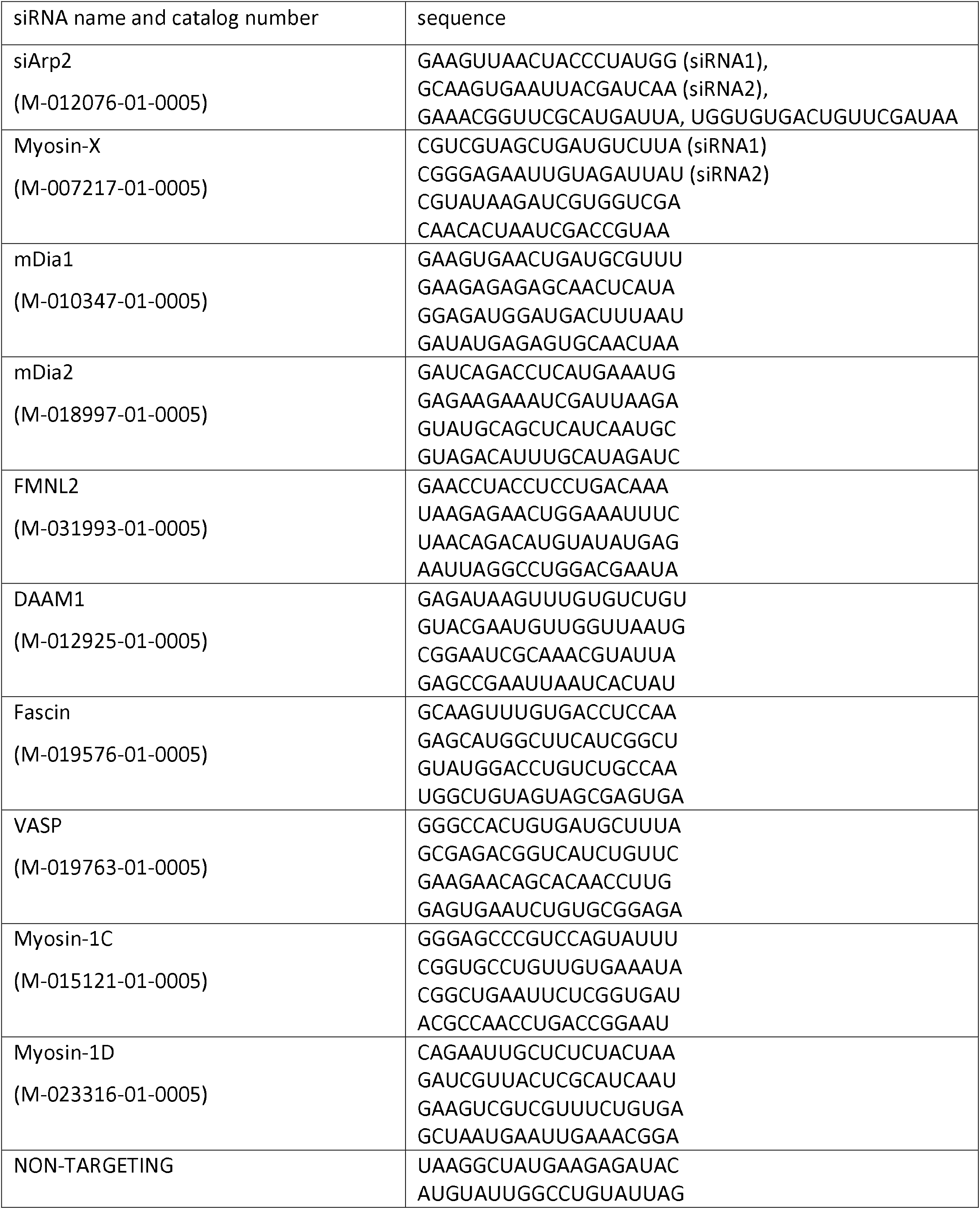
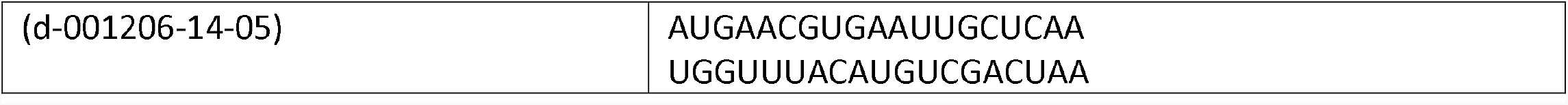
List of siRNAs used.

The degree of protein expression in control and knockdown cells was estimated by western blot. Total cell lysates were prepared using RIPA buffer (Sigma-Aldrich, catalogue no. R0278), supplemented with 10 μL/ml protease inhibitors cocktail (Sigma-Aldrich, catalogue no. P1860). Protein concentration was determined using BCA Protein Assay Kit (Thermo Fisher, 23225) according to manufacturer’ s instructions. Cell lysate was dissolved in Laemmli sample buffer supplemented with 4% 2-mercaptoethanol, separated by 8% SDS-PAGE gels at 100V for 1 hour and then transferred to a 0.45 μm pore size nitrocellulose membrane (Bio-Rad, catalog number: 162-0115) at 0.2 A for 1.5 hours in an ice bath. The nitrocellulose membrane was blocked using 5% BSA in TBS-T (Tris-buffered saline with 0.1% Tween® 20 Detergent) (BioLab Ltd, Israel) for 1 hour at room temperature before incubation at 4 ºC overnight with appropriate primary antibodies diluted in blocking buffer at their respective concentrations summarized in Table 2. After 3 washes in TBS-T, the membrane was probed with the appropriate secondary antibodies conjugated with horseradish peroxidase (Jackson ImmunoResearch Inc, 115-035-003 and 111-035-003) for 1 hour at room temperature. The membrane was then washed for 3 times and developed using SuperSignal ECL reagents (Thermo Fisher Scientific), and visualized using Image lab software.

**Table 2:** List of antibodies used

| Protein | Company | Catalog number | Dilution |
| --- | --- | --- | --- |
| mDia1 | BD Biosciences | 610848 | 1:1000 |
| mDia2 | Santa Cruz Biotechnology Inc | sc-293288 | 1:10 |
| FMNL2 | Santa Cruz Biotechnology Inc | sc-390298 | 1:100 |
| DAAM1 | *Proteintech* | 14876-1-AP | 1:200 |
| Myosin-X | *Sigma-Aldrich* | HPA024223 | 1:500 |
| Arp2 | Abcam | ab49674 | 1:500 |
| VASP | *Weizmann Institute of Science*<br>*bioservices.weizmann.ac.il/antibody/about.html* |  | 1:50 |
| Vinculin | *Weizmann Institute of Science*<br>*bioservices.weizmann.ac.il/antibody/about.html* |  | 1:50 |
| Tubulin | *Sigma-Aldrich* | T6199 | 1:1000 |
| Myosin-1C | Santa Cruz Biotechnology Inc | sc-136544 | 1:200 |
| Myosin-1D | Santa Cruz Biotechnology Inc | sc-515292 | 1:200 |

### Substrate coating

Bacterially expressed recombinant galectin-8 were purified as previously described (Hadari et al., 1995). α-Lactose-Agarose beads used for galectin-8 purification were purchased from Sigma (Catalog no. L7634). The glass-bottomed petri-dishes (MatTek, P35G-1.5-14-C) were incubated with 25 µg/ml galectin-8 solution in PBS for 2 hours at room temperature and washed five times with PBS before the plating of cells.

### Cell suspension preparation

Cells from 70%-80% confluent culture were first washed with warm PBS once, then incubated in Trypsin EDTA Solution B (Trysin 0.25%, EDTA 0.05%) (Biological Industries USA, 03-052-1B) at 37°C for two minutes and gently suspended by addition of serum-free DMEM with trypsin inhibitor (T9003, Sigma-Aldrich) (1mg of trypsin inhibitor per milliliter Trypsin EDTA Solution B). The suspension was centrifuged at 1500 rpm for 5 minutes, supernatant was removed and serum-free medium was added to re-suspend the cells. Then, the cells were plated to the pre-coated petri dish and either imaged or fixed at appropriate time points.

### Drug treatment

For sugar inhibition studies, 10 mM thiodigalactoside (TDG) (17154, Cayman Chemical) was added to cells in suspension and incubated for 10 minutes at 37°C before the cells were seeded on the substrates in the medium containing the same concentration of TDG.

### Immunofluorescence staining

For immunostaining, cells cultured on glass bottomed dish were fixed/permeabilized in phosphate-buffered saline (PBS) containing 0.25% Triton X-100, 0.25% glutaradehyde and 3% paraformaldehyde at 37°C for 15 min. The cells were then twice washed for 10 minutes with PBS. Before staining, the fixed cells were treated with 1mg/ml sodium borohydride in PBS for 15 min on ice. The cells were then washed with PBS, incubated with blocking solution (5% bovine serum albumin (BSA) in PBS) for 1 h at room temperature and washed with PBS again. The cells were incubated with appropriate primary antibodies at room temperature for one hour, washed three times with PBS, and then incubated with appropriate fluorescently labelled secondary antibodies or/and phalloidin to visualize actin. Goat anti-Rabbit/mouse IgG (H+L) Cross-Adsorbed ReadyProbes™ Secondary Antibody, Alexa Fluor 488/647 was purchased from Thermo Fisher (catalogue no. R37116/ R37114, A21245/A32728), and was used at dilution 1:400. Phalloidin–Tetramethylrhodamine B isothiocyanate was obtained from Sigma-Aldrich (catalogue no. P1951) and was used at 1:400 dilution.

### Microscopy and live cell imaging

Cells were plated at a density of 5 × 10^4^ cells ml^−1^ onto the 35 mm cell culture dish with 14 mm-diameter glass bottom (MatTek, catalogue number: P35G-1.5-14-C) coated with galectin-8 as described above. Video recordings started 5 minutes following cell seeding. Interference reflection microscopy (IRM) time-lapse imaging were carried out using the DeltaVision RT microscopy system (Applied Precision Inc., Issaquah, WA, USA), equipped with a ×100 oil immersion objective (1.3 NA, UPlanSApo), at 5 seconds time intervals between frames. Confocal images and videos were taken with ANDOR Dragonfly spinning disk confocal microscope using 100X objective and a sCMOS (Zyla) camera. Super-resolution SIM imaging was performed using W1-spinning-disc confocal unit coupled with the live super-resolution (SR) module (instant SIM) (GatacaSystems), mounted on Eclipse microscope (Nikon) with Perfect Focus System, supplemented with the objective Plan Apo 100x oil NA1.45 and scientific complementary metal–oxide–semiconductor (sCMOS) camera Prime95B (Photometrics).

### Cryo-electron tomography

Cells were applied onto galectin-8 coated EM grids with carbon support film (R2/2, Au mesh; Quantifoil, Jena, Germany). After 15-20 min incubation, a 4µl drop of fiducial gold marker (10nm; Aurion, Wageningen, Netherlands) was added to the sample prior to plunge freezing into liquid ethane. A Titan Krios transmission electron microscope (Thermo Fisher Scientific, Waltham, USA) equipped with a Quantum energy filter and a K2-Summit direct electron detector (Gatan, Pleasanton, USA) was used for cryo-EM data acquisition.

The tomograms were recorded with an electron flux of around 10 electrons per pixel per second using SerialEM (Mastronarde, 2005). Tilt series were acquired at a magnification of 64’ 000x with exposure time 1.6 sec, and a dose-fractionated frame rate at 6.25 frame/sec. The tilt-series covered an angular range of -60° to +60°, and were recorded with tilt increments of 3° at a defocus of -4⍰μm. The accumulated electron dose was around 134 e-/Å2. Finally, the tomograms were reconstructed in IMOD (Kremer et al., 1996) and the isosurface visualizations were rendered with AMIRA (Thermo Fisher Scientific, Waltham, USA).

### Filopodia image analysis

Filopodia length and number were measured automatically using Matlab programs from confocal images of cells stained with phalloidin, as described previously (Li et al., 2021b). Briefly, intensity thresholding by Otsu’ s method (Otsu, 1979) was firstly applied to segment the cell. Then filopodia were identified using FiloDetect algorithm (Nilufar et al., 2013). Filopodia were defined as high aspect ratio (≥1.5: 1) objects protruding from the “main cell body” with a smooth boundary. The filopodia shorter than 0.6 µm were ignored.

For chirality measurement, all images were arranged as if the observer looked at the cells from the medium towards the substrate. This was routinely checked by filming of asymmetric test-object in the same way as the cells. The assessment of left-bent, right-bent and wavy filopodia was performed manually in a blind fashion (without the knowledge of the condition of the experiments).

Box and whisker plots show median values (middle line inside the box), mean (circle), upper and lower quartiles (upper and lower bound of the box), and outliers (whiskers) (values which are 1.5 times larger than the upper or 1.5 times smaller than the lower quartiles). In chirality bar plots, the height of the bar indicates the mean, and error bar indicates the standard deviation. Calculation of p value was performed using two-tailed Mann-Whitney tests using OriginLab software.

